# Genetic evidence for an origin of the Armenians from Bronze Age mixing of multiple populations

**DOI:** 10.1101/015396

**Authors:** Marc Haber, Massimo Mezzavilla, Yali Xue, David Comas, Paolo Gasparini, Pierre Zalloua, Chris Tyler-Smith

**Affiliations:** The Wellcome Trust Sanger Institute, Wellcome Trust Genome Campus, Hinxton, Cambs. CB10 1SA, United Kingdom.; Institute for Maternal and Child Health -IRCCS “Burlo Garofolo” - Trieste, University of Trieste, Italy.; Institut de Biologia Evolutiva (CSIC–UPF), Departament de Ciències de la Salut i de la Vida, Universitat Pompeu Fabra, Barcelona, Spain; The Lebanese American University, Chouran, Beirut, Lebanon

## Abstract

The Armenians are a culturally isolated population who historically inhabited a region in the Near East bounded by the Mediterranean and Black seas and the Caucasus, but remain underrepresented in genetic studies and have a complex history including a major geographic displacement during World War One. Here, we analyse genome-wide variation in 173 Armenians and compare them to 78 other worldwide populations. We find that Armenians form a distinctive cluster linking the Near East, Europe, and the Caucasus. We show that Armenian diversity can be explained by several mixtures of Eurasian populations that occurred between ∼3,000 and ∼2,000 BCE, a period characterized by major population migrations after the domestication of the horse, appearance of chariots, and the rise of advanced civilizations in the Near East. However, genetic signals of population mixture cease after ∼1,200 BCE when Bronze Age civilizations in the Eastern Mediterranean world suddenly and violently collapsed. Armenians have since remained isolated and genetic structure within the population developed ∼500 years ago when Armenia was divided between the Ottomans and the Safavid Empire in Iran. Finally, we show that Armenians have higher genetic affinity to Neolithic Europeans than other present-day Near Easterners, and that 29% of the Armenian ancestry may originate from an ancestral population best represented by Neolithic Europeans.

## Introduction

Insights into the human past come from diverse areas including history, archaeology, linguistics and, increasingly, genetics. Observed patterns of present-day genetic diversity can be compared with models that include past population processes such as migration, divergence and admixture, and the best model chosen. These models often require representing ancestral populations and mostly consider present-day populations as direct descendants of the ancient inhabitants of a region. However, archaeological and genetic data reveal that human history has often been shaped by regional or localized population movements that can confound simple demographic models.^1,2^ Ancient DNA (aDNA) studies have also shown that the genetic landscape has been continuously shifting,^3,4^ possibly triggered by environmental and cultural transitions. aDNA research is useful for understanding past demographic events; however, samples are limited and obtaining aDNA from warm climates remains a challenge. We have previously shown that studying genetic isolates also provides insights into human genetic variation and past demographic events.^5^ For example, by studying Jews, Druze, and Christians from the Near East, we showed the region had more genetic affinity to Europe 2,000 years ago than at present.^5^

In the present study, we investigate the Armenians, a population today confined to the Caucasus but who occupied Eastern Anatolia, reaching as far as the Mediterranean coast, up until the start of the 20^th^ century (Figure 1). Political turmoil in the region during World War One resulted in the displacement of the Armenian population and its restriction today to an area in the Caucasus between the Black and Caspian seas. Armenians are an ethno-linguistic-religious group distinct from their surrounding populations. They have their own church, the Armenian Apostolic Church, which was founded in the 1st century CE, and became in 301 CE the first branch of Christianity to become a state religion. They have also their own alphabet and language which is classified as an independent branch of the Indo-European language family. The Armenian language is a subject of interest and debate among linguists for its distinctive phonological developments within Indo-European languages and for its affinity to Balkan languages such as Greek and Albanian. The historical homeland of the Armenians sits north of the Fertile Crescent, a region of substantial importance to modern human evolution. Genetic and archaeological data suggest farmers expanding from this region during the Neolithic populated Europe and interacted/admixed with pre-existing hunter-gatherer populations.^6^ Furthermore, Armenia’s location may have been important for the spread of Indo-European languages, since it is believed to encompass or be close to the Proto-Indo-European homeland (Anatolia or Pontic Steppe) from which the Indo-Europeans and their culture spread to Western Europe, Central Asia and India.

**Figure 1.**
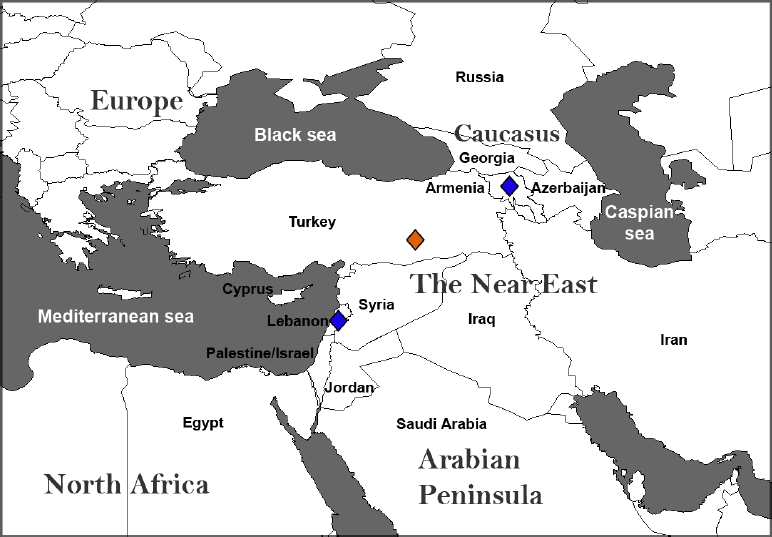
Map of the Near East and surrounding regions. The map shows location of present-day Armenia and neighbouring countries. Blue lozenges show the recruitment sites for Armenian samples used in this study. Political turmoil during World War One resulted in the displacement of the East Turkey Armenian population (orange lozenge) to present-day Armenia or to several other nearby countries such as Lebanon.

Previous genetic studies on Armenians are scarce and genome-wide analysis is limited to a few Armenian samples in broad surveys without any detailed analysis. Armenians were found to have genetic affinity to populations including the Jews, Druze and Lebanese Christians, in addition to showing genetic continuity with the Caucasus.^5,7,8^

In this study, we analyse newly-generated genome-wide data from Armenians as well as individuals from 78 other worldwide populations. We investigate genetic signatures of past events such as the emergence of Armenians as an ethnic group, cultural changes in the Near East, and the expansions of ancient populations in this region.

## Materials and Methods

### Subjects and the genetic datasets

Armenian samples were collected from Lebanon (n=39) and Armenia: Chambarak (n=30), Deprabak (n=18), Gavar (n=12), Martumi (n=19), Yegvard (n=11), and Yerevan (n=9). Armenian individuals recruited in Lebanon traced their ancestry to East Turkey; they signed informed consent approved by the IRB of the Lebanese American University and were genotyped on Illumina 610K or 660K bead arrays.

Armenian subjects recruited from present-day republic of Armenia signed consents approved by the ethical committee of the Maternal and Child Health Institute IRCCS-Burlo Garofolo Hospital (Trieste, Italy). Samples were genotyped on Illumina HumanOmniExpress and described previously by Mezzavilla et al.^9^

Genotype data can be downloaded from: ftp://ngs.sanger.ac.uk/scratch/project/team19/Armenian Additional Armenian samples (n=35) were added along with 1,509 samples from the literature that represent 78 worldwide populations.^5,7,8,10^

PLINK^11^ was used for data management and quality control. The required genotyping success rate was set to 99%, sex-linked and mitochondrial SNPs removed, leaving 300,899 SNPs.

### Population Structure

Principal components were computed with EIGENSOFT v 4.2^12^ using 78 global populations, and the Armenian samples were projected onto the plot. The Bayesian Information Criterion (BIC) was computed by mclust (http://www.stat.washington.edu/mclust) over the first 10 principal components of the projected Armenian samples on the global PCA. The best model to classify the Armenians according to the BIC values is with three components (clusters) (Supplementary Figure S1).

Inference of population relations from haplotypes was assessed using Chromopainter^13^ with 10,000,000 burnin and runtime and 10,000 MCMC samples. A bifurcating tree of relationships amongst these populations was built using fineSTRUCTURE^13^ (Supplementary Figure S2).

Effective population size was estimated from linkage disequilibrium and time of divergence between populations was calculated using *NeON*^14^ with default parameters. Genetic distance (*Fst*) between populations was calculated using the software 4P^15^. The generation time used was 28 years.

### Admixture analysis

We used *f3* statistics^16^ *f3(A; B,C)* where a significantly negative statistic provides evidence that *A* is derived from admixture of populations related to *B* and *C*. We tested all possible *f3* statistics in our dataset and calculated standard errors using blocks of 500 SNPs.^17^ To date the time of admixture, we used *ALDER*^18^ which computes the weighted LD statistic to make inferences about population admixture. The reference populations consisted of 1300 samples and 53 populations reduced from the original dataset after removing populations that are themselves highly admixed (Supplementary Table 1). We collected results that were significant (z-score >|4|) and summarized the findings in Table 1 after pooling populations into respective geographical groups. Sardinians appear to have a distinctive admixture pattern from other West Europeans and therefore are shown separately.

**Table 1.** Source populations and admixture time for Armenians

| **Source 1** | **Source 2** | **f3-statistics†** | **z-score** | **Time ± se** | **p-value** |
| --- | --- | --- | --- | --- | --- |
| West Europeans | Sub-Saharan Africans | -0.00105083 | -5.01009 | 5826.52 ± 672.84 | 2.00E-08 |
| Caucasus populations | Sardinians | -0.000251561 | -4.89462 | 5554.64 ± 361.76 | 4.10E-36 |
| Central and South Asians | Sardinians | -0.00110183 | -16.261 | 4886.28 ± 271.32 | 3.00E-34 |
| Caucasus populations | Arabian Peninsula populations | -0.000467324 | -7.32053 | 4812.08 ± 381.64 | 8.10E-22 |
| Caucasus populations | North Levantines | -0.000237086 | -5.99602 | 4673.2 ± 343 | 9.10E-19 |
| South Levantines | Caucasus populations | -0.000235109 | -5.03679 | 4376.12 ± 426.16 | 7.60E-19 |
| Central and South Asians | West Europeans | -0.000345672 | -5.42631 | 4233.04 ± 1111.6 | 0.0029 |
| North Europeans | Arabian Peninsula populations | -0.000433178 | -5.1784 | 3939.88 ± 239.96 | 1.70E-32 |
| Central and South Asians | North Levantines | -0.000484051 | -6.95695 | 3908.52 ± 682.08 | 2.10E-07 |

† lowest f3 result from each source region

For tests of genetic affinity to Neolithic Europeans, we merged our samples with the genome of the Tyrolean Iceman.^19^ We downloaded the BAM file mapped to hg18 and called all variants using *GATK*^20^. liftOver (http://genome.ucsc.edu) was used to convert the coordinates to hg19. The final dataset consisted of 91,115 SNPs.

For tests of genetic affinity to Mesolithic Europeans, we merged our dataset with the genome of the La Braña sample.^21^ We downloaded the BAM file mapped to hg19 and called the variants using *GATK*. The final dataset consisted of 103,627 SNPs.

We applied *TreeMix*^17^ rooting the tree with a Denisovan genome, and standard errors were estimated using blocks of 500 SNPs. We generated 100 bootstrap replicates by resampling blocks of 500 SNPs to assess the stability of the tree topology. We used outgroup *f3* statistics^3,22^ in the form of *f3(Yoruba; Iceman, X)* and *f3(Yoruba; La Braña, X)* to assess the shared genetic history of the ancient Europeans with the modern populations. In the absence of admixture with Yoruba, deviation from 0 will be a function of the shared genetic history of the ancient Europeans and the non-African population.

## Results

### Armenians’ relationship to world populations

To study Armenians’ genetic relationship to worldwide populations, we computed principal components using 78 populations (Supplementary Table 1) and projected the Armenians onto the plot in a procedure called “PCA projection”^23^ (Figure 2a) which ensures that the PCA patterns are not affected by the large number of Armenians used in the analysis. We observe that Armenians form a distinctive cluster bounded by Europeans, Near Easterners, and the Caucasus populations. More specifically, Armenians are close to 1) Spaniards, Italians and Romanians from Europe; 2) Lebanese, Jews, Druze and Cypriots from the Near East; and 3) Georgians and Abkhazians from the Caucasus (Figure 2b). The position of the Armenians within the global genetic diversity is unique and appears to mirror the geographical location of Anatolia. Previous genetic studies have generally used Turks as representatives of ancient Anatolians. Our results show that Turks are genetically shifted towards Central Asians, a pattern consistent with a history of mixture with populations from this region.

**Figure 2.**
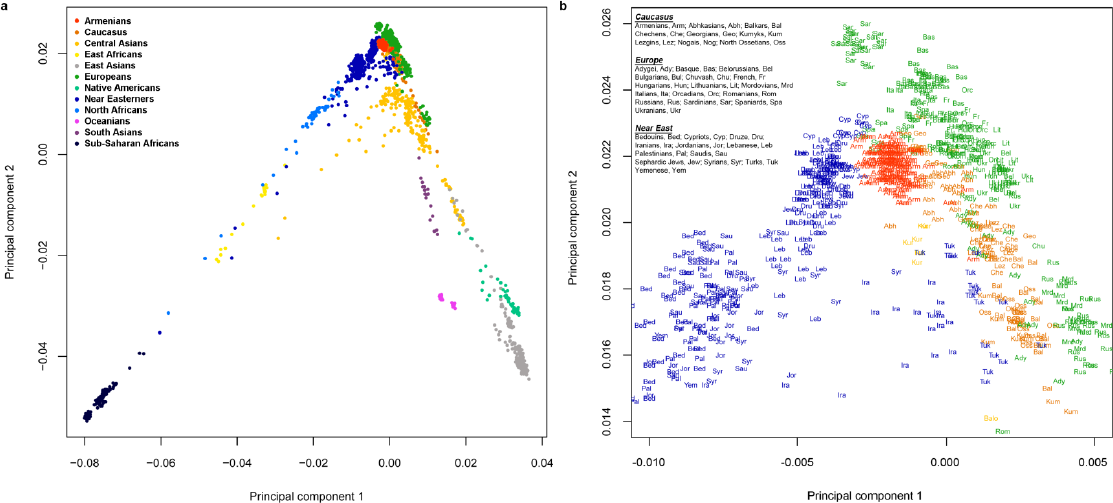
Principal component analysis of >240,000 SNPs showing the top two components. a) The position of Armenians in a global genetic diversity sample based on 78 populations from 11 geographical regions. Armenians (173 individuals) were projected to the plot and therefore did not contribute to the observed global structure. b) A magnification shows that the Armenians (red) demonstrate genetic continuity with the Near East, Europe, and the Caucasus.

These diversity patterns observed in the PCA motivated formal testing of admixture in Armenians and other regional populations.

### Admixture in the Near East

To formally test for population mixture in Armenians we performed a *3-population* test^24^ in the form of *f3(Armenian; A, B)* where a significantly negative value of the *f3* statistic implies that Armenians descend from a mixture of the populations represented by *A* and *B*, chosen from the 78 global populations. We found signals of mixture from several African and Eurasian populations (Table 1, Figure 3). The most significantly negative *f3* statistics are for mixture of populations related to Sardinians and Central Asians, followed by several mixtures of populations from the Caucasus, Arabian Peninsula, the Levant, Europe, and Africa. We sought to date these mixture events using exponential decay of admixture-induced linkage disequilibrium (LD). The oldest mixture events appear to be between populations related to sub-Saharan Africans and West Europeans occurring ∼3,800 BCE, followed closely by mixture of Sardinian and Caucasus-related populations. Later, several mixture events occurred from 3,000–1,200 BCE involving diverse Eurasian populations (Table 1, Figure 3). We compared patterns of admixture in Armenians to other regional populations and detected signals of recent admixture in most other populations. For example, we find 7.9% (±0.4) East Asian ancestry in Turks from admixture occurring 800 (±170). We also detect sub-Saharan African gene flow 850 (±85) years ago in Syrians, Palestinians and Jordanians.

**Figure 3.**
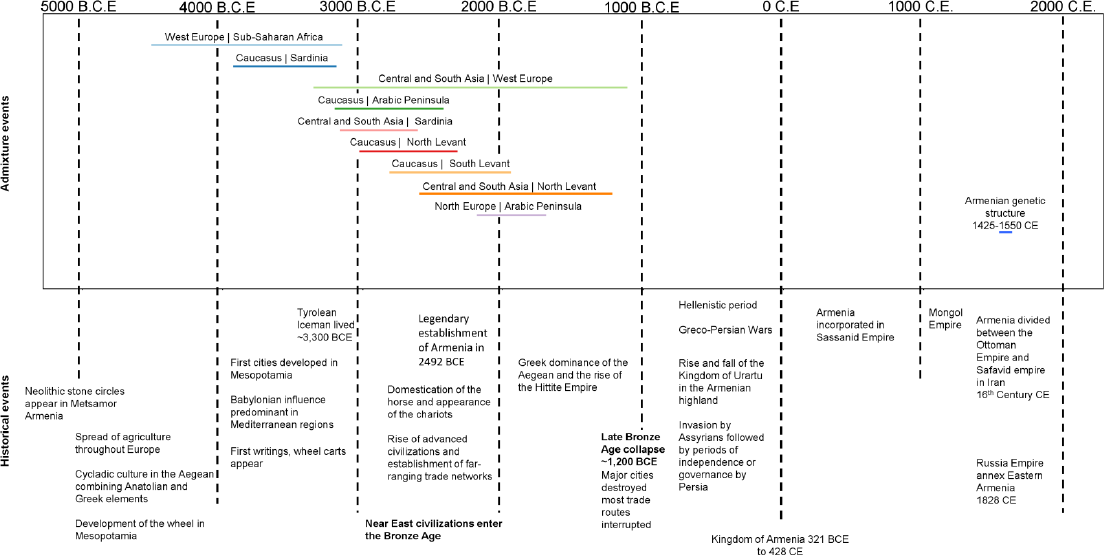
Genetically-inferred source populations for Armenians, admixture times and genetic structure. Admixture events were estimated using decay of linkage disequilibrium with regional populations as sources for Armenians. Each horizontal coloured line indicates an admixture event and its width reflects the estimated date of admixture and standard error. The plot also shows the estimated date of establishment of genetic structure within Armenians (1425-1550 CE). Major historical events and cultural developments in the Near East are at the bottom.

### Structure of the Armenian population

To investigate the presence of genetic structure within the Armenian population, we performed model-based clustering on the values of the Armenian samples from the global PCA. We observe the following: 1) Armenians in the diaspora that trace their origin to historical Western Armenia (modern-day East Turkey) form one group (Supplementary Figure S1, Cluster 1). 2) Armenians in modern-day Armenia (historical Eastern Armenia) are split in two major groups: 33% are in Cluster 1 and 57% form Cluster 2 (Supplementary Figure S1). This structure could be the result of the Western Armenians’ migration to the East after the events of 1915 CE that displaced the entire Western Armenian population. 3) A few Armenians recruited from Chambarak and Maykop (Republic of Adygea, Russia), form an outlier to the two major Armenian clusters (Supplementary Figure S1, Cluster 3).

We investigated Armenian structure further using a procedure called “chromosome painting”^13^ which reconstructs the haplotype of every individual (receiver) in our dataset using the haplotypes of other individuals (donors) in the dataset. We then constructed a tree that infers population relationships and similarities (Supplementary Figure S2). We found, similarly to our previous clustering results, fine genetic structure that splits Armenians into two major groups that are more similar to each other than to any other global population. We estimate from the LD patterns that divergence between the two major Armenian groups started 450–575 years ago (Figure 3).

### Relationship to ancient Europeans

We merged our dataset with the genome of the Tyrolean Iceman, a 5,300-year-old individual discovered on the Italian part of the Ötztal Alps. We used *TreeMix*^17^ to construct a tree of genetic relationships using representative regional populations plus Armenians and Turks from the Near East. *TreeMix* uses a model that allows for both population splits and gene flow to better capture historical relationships between populations. We obtained a tree that recapitulates the known relationships among population groups. Furthermore, the tree shows the Iceman shared drift with Sardinians, as previously reported.^19^ We then ran *TreeMix* allowing it to infer only one migration event, and revealed gene flow from the Iceman to Armenians accounting for about 29% of their ancestry. The graph structure appeared robust in 100 bootstrap replicates with the first migration (highest weight and lowest p-value) always leading from the Iceman to Armenians (Figure 4).

**Figure 4.**
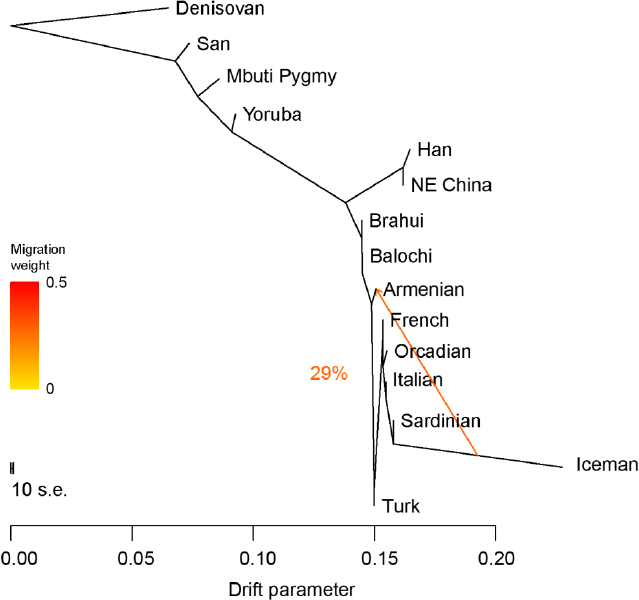
Inferred population tree with one mixture event. The graph was inferred by *TreeMix* allowing one migration event. The migration arrow is coloured according to its weight; the weight is correlated with the ancestry fraction and shows that 29% of Armenian ancestry is derived from a population related to ancient Europeans. The graph is stable in 100 bootstrap replicates.

This structure was further investigated using outgroup *f3* statistics.^3,22^ The expected value of *f3(Yoruba; Iceman, X)* in the absence of admixture with Yoruba, will be a function of the shared genetic history of the Iceman and *X* (non-African populations). Most shared ancestry with the Iceman is with Sardinians and other Europeans (Supplementary Figure S3). This is directly followed by shared ancestry with some Near Eastern populations: Cypriots, Sephardic Jews, Armenians, and Lebanese Christians. Other Near Easterners such as Turks, Syrians, and Palestinians show less shared ancestry with the Iceman.

To investigate if the affinity of the Near East genetic isolates to Europeans precedes the arrival of the early farmers to Europe (represented by the Iceman), we repeated the outgroup *f3* statistics and replaced the Iceman with a 7,000 year old European hunter gatherer from Spain (La Braña).^21^ West European hunter-gatherers have previously been shown to have contributed ancestry to all Europeans but not to Near Easterners.^6^ Consistent with this, we found reduced affinity and no noticeable structure in the Near Easterners in their relation to La Braña (Supplementary Figure S4) (compared with the Iceman).

## Discussion

The origins of the Armenians and their cultural uniqueness are poorly understood. Here, we investigate the information that can be obtained by genetic analysis of present-day Armenians and comparisons with other present-day and ancient samples.

The position of the Armenians within global genetic diversity is unique and appears to mirror the geographical location of Anatolia, which forms a bridge connecting Europe, the Near East and the Caucasus. Anatolia’s location and history have placed it at the centre of several modern human expansions in Eurasia: it has been inhabited continuously since at least the early Upper Palaeolithic,^25^ and has the oldest known monumental complex built by hunter-gatherers in the 10th millennium BCE.^26^ It is believed to have been the origin and/or route for migrating Near Eastern farmers towards Europe during the Neolithic,^27^ and has also played a major role in the dispersal of the Indo-European languages.^28^

We investigated Armenians further by inferring their admixture history. The Armenians show signatures of an origin from mixture of diverse populations occurring 3,000 to 2,000 BCE. This period spans the Bronze Age, characterized by extensive use of metals in farming tools, chariots and weapons, accompanied by development of the earliest writing systems and the establishment of trade routes and commerce. Many civilizations such as in Ancient Egypt, Mesopotamia, and the Indus valley grew to prominence. Major population expansions followed, triggered by advances in transportation technology and the pursuit of resources. Our admixture tests show that Armenians genomes carry signals of extensive population mixture during this period. We note that these mixture dates also coincide with the legendary establishment of Armenia in 2,492 BCE. Admixture signals decrease to insignificant levels after 1,200 BCE, a time when Bronze Age civilizations in the eastern Mediterranean world suddenly collapsed, with major cities being destroyed or abandoned and most trade routes disrupted. This appears to have caused Armenians’ isolation from their surroundings, subsequently sustained by the cultural/linguistic/religious distinctiveness that persists until today. The genetic landscape in most of the Middle East appears to have been continuously changing since then. For example, we detect East Asian ancestry in Turks from admixture occurring 800 (±170) years ago coinciding with the arrival of the Seljuk Turks in Anatolia from their homelands near the Aral Sea. We also detect sub-Saharan African gene flow 850 (±85) years ago in Syrians, Palestinians and Jordanians, consistent with previous reports on recent gene flow from Africans to Levantine populations after the Arab expansions.^5,29^ The admixture pattern in Armenians appears similar to patterns we have observed in some other genetic isolates in the region, such as Sephardic Jews and Lebanese Christians, who show limited admixture with culturally different neighbouring populations in the last two millennia.^5^ Our tests suggest that Armenians had no significant mixture with other populations in their recent history and have thus been genetically isolated since the end of the Bronze Age, 3,000 years ago. In recent times, we detect a genetic structure within the Armenian population that developed ∼500 years ago. The date coincides with the start of the Ottoman-Persian wars and the split of Armenia into West and East between the Ottoman Empire in Turkey and the Safavid Empire in Iran.

One of the most-studied demographic processes in population genetics is the Neolithic expansion of Near Eastern farmers into Europe beginning ∼8,000 years ago. Armenians’ location at the northern tip of the Near East suggests a possible relationship to the expanding Neolithic farmers. We find in Armenians and other genetic isolates in the Near East high shared ancestry with ancient European farmers with ancestry proportions similar to present-day Europeans but not to present-day Near Easterners. These results suggest that genetic isolates in the Near East - Cypriots (an island population), Near Eastern Jews and Christians (religious isolates), and Armenians (Ethno-linguistic isolate) - probably retain features of an ancient genetic landscape in the Near East that had more affinity to Europe than the present populations do. Our tests show that most of the Near East genetic isolates ancestry shared with Europeans can be attributed to expansion after the Neolithic period.

Armenians’ adoption of a distinctive culture early in their history resulted in their genetic isolation from their surroundings. Their genetic resemblance today to other genetic isolates in the Near East, but not to most other Near Easterners, suggests that recent admixture has changed the genetic landscape in most populations in the region. Armenians’ genetic diversity reveals that the ancient Near East had higher affinity to Neolithic Europe than it does now, and that Bronze Age demographic processes had major impact on the genetics of populations in this region.

The importance of populations like the Armenians is not limited to the study of past demographic processes; isolated populations are emerging as a powerful tool for many different genetic investigations such as rare variant associations with complex phenotypes and the characterization of gene-environment interactions.^30^ Armenians emergence from founders in the Bronze Age, accompanied by a long period of subsequent isolation, may have enriched rare disease alleles and therefore merits future medical exploration.

## Acknowledgments

We thank Qasim Ayub for his comments and suggestions on the manuscript. We also thank Collin Renfrew and Merritt Ruhlen for their comments at early stages of this study. This work was supported by Wellcome Trust grant 098051.

## Supplementary info

**Figure S1.**
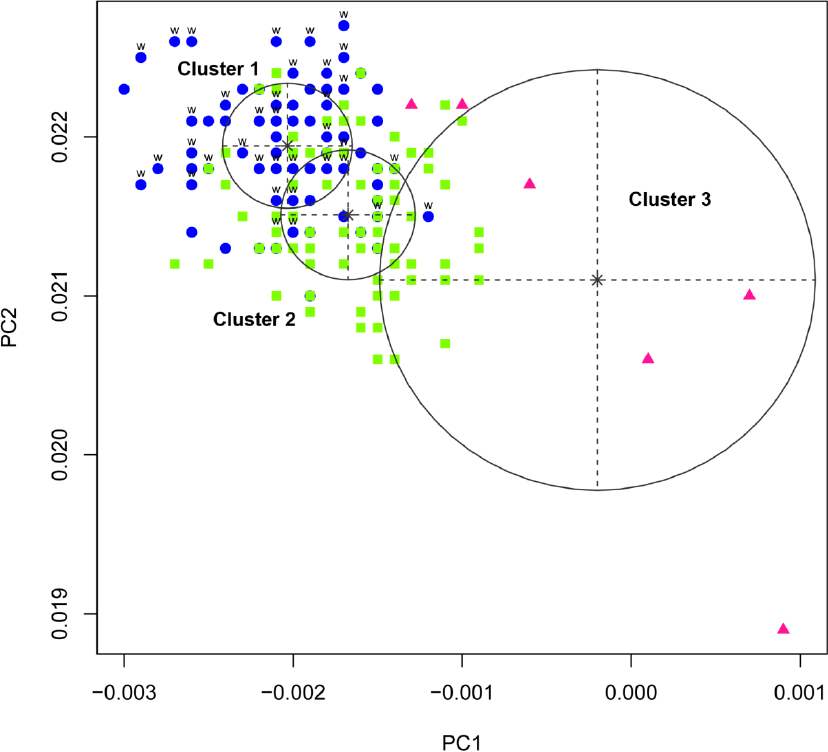
Genetic structure in Armenians. *MCLUST* classifies Armenians using a Bayesian Information Criterion into three clusters. Cluster 1 (blue) includes 95% of the Armenians that trace their origin to Western Armenia (East Turkey) (labelled W). Cluster 1 also includes 33% of the general Armenians (recruited from modern Armenia). Cluster 2 includes 57% of the general Armenians. Cluster 3 includes six Armenians recruited from Chambarak (modern-day Armenia) or Maykop (Republic of Adygea, Russia).

**Figure S2.**
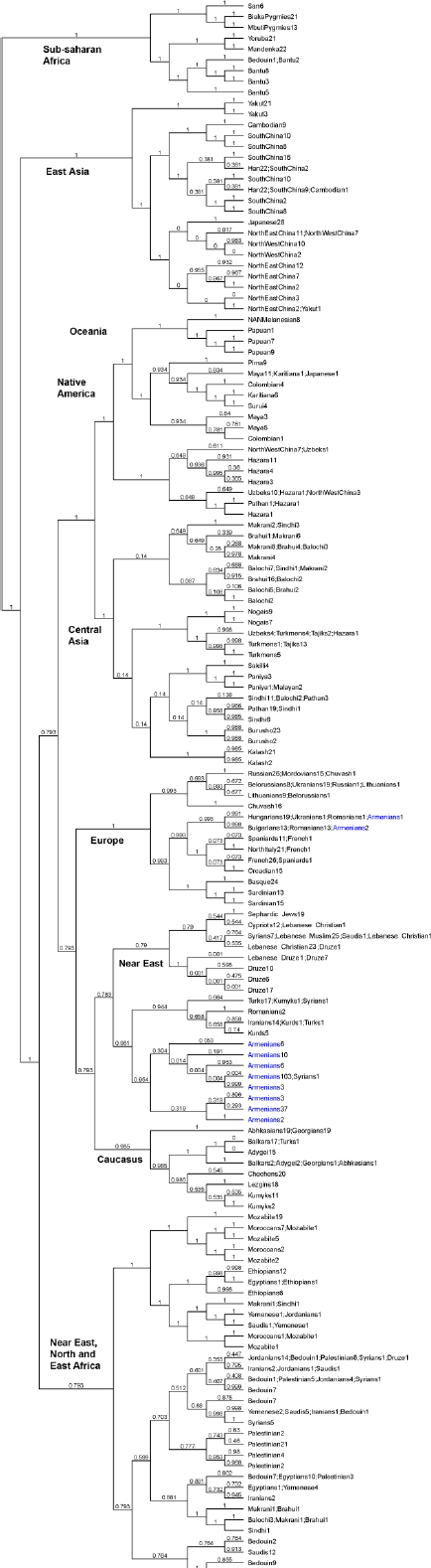
Population relationships from genome-wide haplotypes. Each tip of the tree corresponds to an individual; numbers of individuals are shown next to their population name at the tip of the branches. Numbers on branches show partition posterior probability. Armenians are shown in blue, forming two major clusters in a Near Eastern branch.

**Figure S3.**
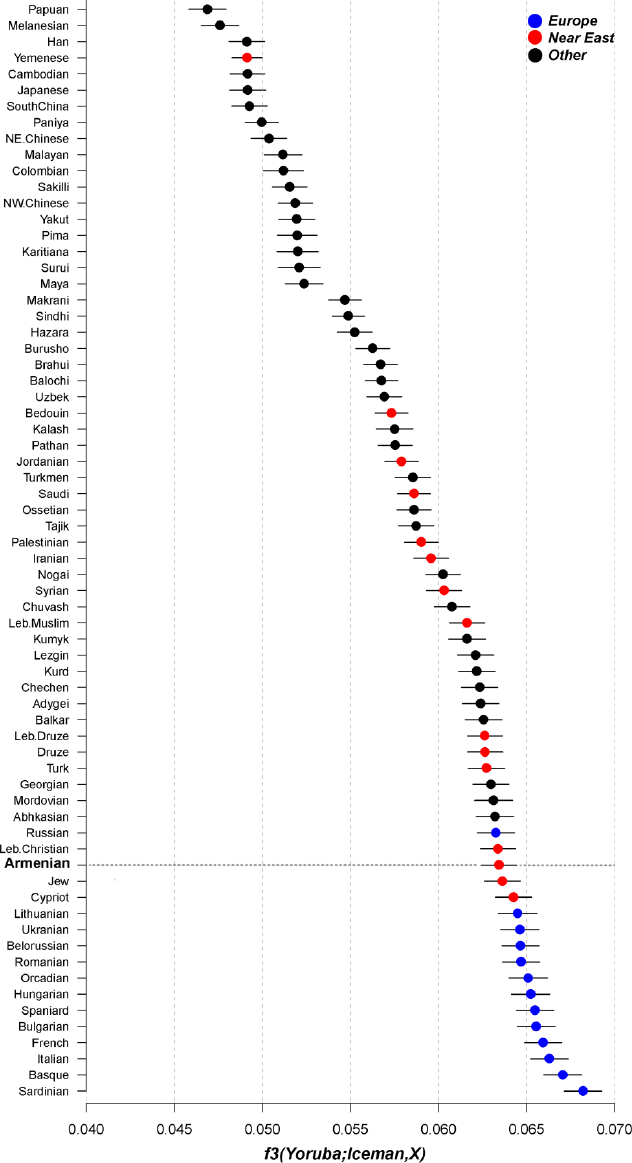
Shared genetic drift between worldwide populations and the Tylorean Iceman, a 5,300 year old European.

**Figure S4.**
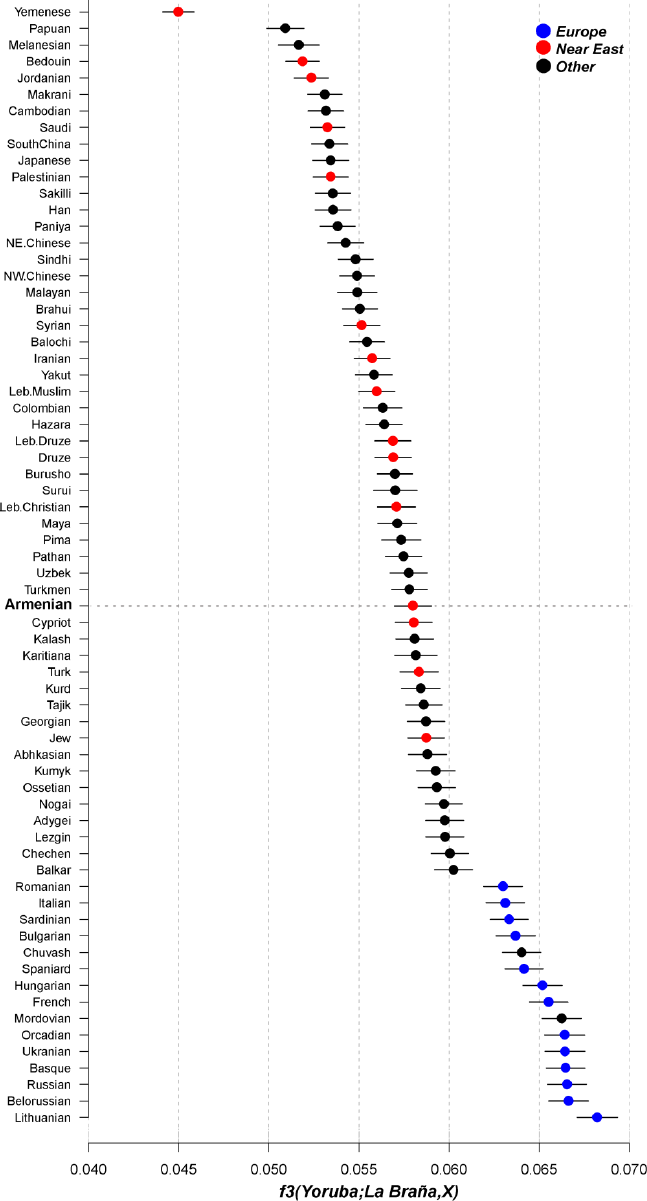
Shared genetic drift between worldwide populations and La Braña, a 7,000 year old European.

**Table S1.**
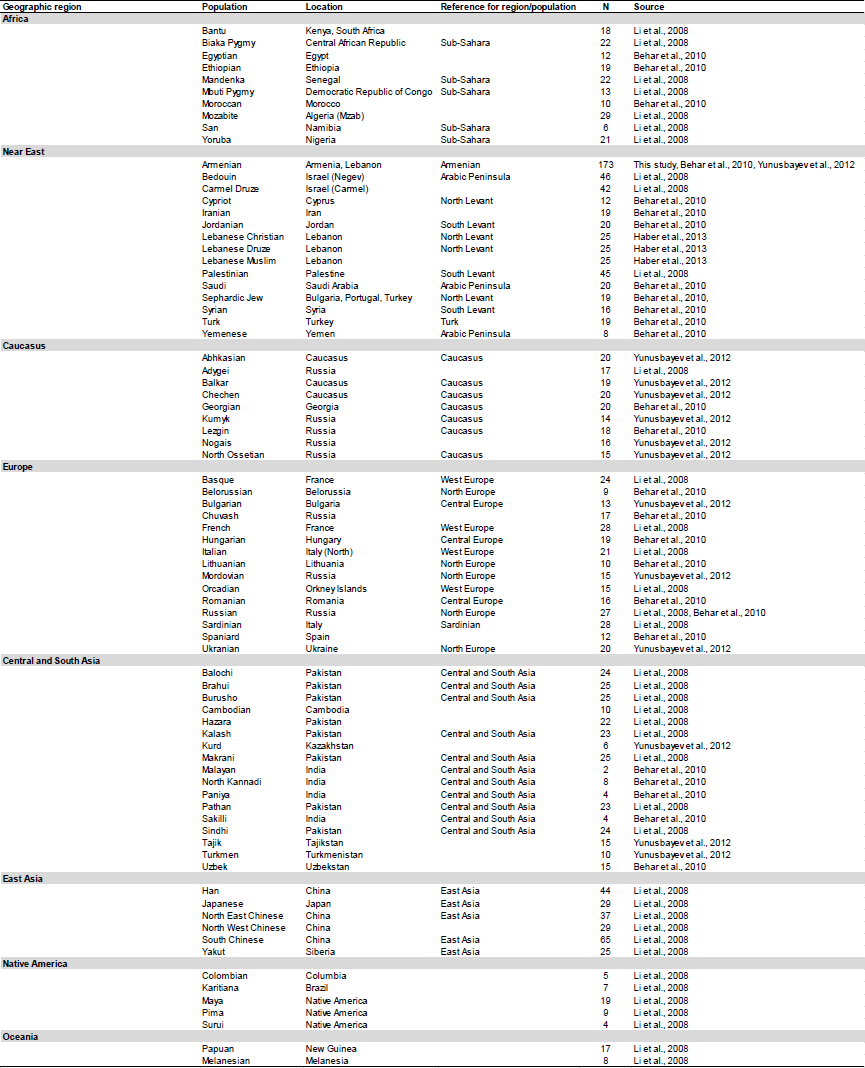
Populations selected for this study

